# Genetic analysis of retinal cell types reveals synaptic pathology in schizophrenia

**DOI:** 10.1101/2024.08.09.607343

**Authors:** Emanuel Boudriot, Marius Stephan, Finn Rabe, Lukasz Smigielski, Andrea Schmitt, Peter Falkai, Michael J. Ziller, Moritz J. Rossner, Philipp Homan, Sergi Papiol, Florian J. Raabe

**Affiliations:** Max Planck Institute of Psychiatry, 80804 Munich, Germany; Department of Psychiatry and Psychotherapy, LMU University Hospital, LMU Munich, 80336 Munich, Germany; Systasy Bioscience GmbH, 81669 Munich, Germany; Department of Adult Psychiatry and Psychotherapy, University of Zurich, 8032 Zurich, Switzerland; Department of Child and Adolescent Psychiatry and Psychotherapy, University Hospital of Psychiatry Zurich, University of Zurich, 8032 Zurich, Switzerland; 6German Center for Mental Health (DZPG), partner site Munich-Augsburg; Laboratory of Neurosciences (LIM-27), Institute of Psychiatry, University of São Paulo (USP), São Paulo-SP 05403-903, Brazil; Department of Psychiatry, University of Münster, 48149 Münster, Germany; Center for Soft Nanoscience, University of Münster, 48149 Münster, Germany; Neuroscience Center Zurich, University of Zurich and ETH Zurich, 8057 Zurich, Switzerland; Institute of Psychiatric Phenomics and Genomics (IPPG), LMU Munich, 80336 Munich, Germany

**Author notes:** Corresponding author: Florian J. Raabe, MD, PhD. These authors contributed equally (Emanuel Boudriot, Marius Stephan, Finn Rabe).

## Abstract

**Importance:** As an accessible part of the central nervous system, the retina provides a unique window to study pathophysiological mechanisms of brain disorders in humans. Imaging and electrophysiological studies have revealed retinal alterations across several neuropsychiatric and neurological disorders. However, it remains largely unclear whether primary disease mechanisms within the retina contribute to the observed retinal alterations and which specific retinal cell types and biological mechanisms are involved.

**Objective:** To determine whether specific retinal cell types are affected by genomic risk for neuropsychiatric and neurological disorders and to explore the mechanisms through which genomic risk converges in these cell types.

**Design, Setting, and Participants:** In this study, we combined findings from genome-wide association studies in schizophrenia, bipolar disorder, major depressive disorder, multiple sclerosis, Parkinson disease, Alzheimer disease, and stroke with retinal single-cell transcriptomic data sets from humans, macaques, and mice. To identify susceptible cell types, we applied MAGMA cell type enrichment analyses and performed subsequent pathway analyses. Furthermore, we translated the cellular top hit to the structural level by using retinal optical coherence tomography and genotyping data in the large population-based UK Biobank cohort (*n* = 36,349).

**Main Outcomes and Measures:** Cell type-specific enrichment of genetic risk loading for neuropsychiatric and neurological disorder traits in the gene expression profiles of retinal cells.

**Results:** Amacrine cells (interneurons within the retina) were robustly enriched in schizophrenia genetic risk across mammalian species and in different developmental stages. This enrichment was primarily driven by genes involved in synapse biology. On the structural level, higher polygenic risk for schizophrenia was associated with thinning of the ganglion cell–inner plexiform layer, which contains dendrites and synaptic connections of amacrine cells. Moreover, retinal immune cell populations were enriched in multiple sclerosis genetic risk. No consistent cell type associations were found for bipolar disorder, major depressive disorder, Parkinson and Alzheimer disease, or stroke.

**Conclusions and Relevance:** This study provides novel insights into the cellular underpinnings of retinal alterations in neuropsychiatric and neurological disorders and highlights the retina as a potential proxy to study synaptic pathology in schizophrenia.

## Introduction

One major challenge in neuropsychiatric research is the difficulty of accessing human brain tissue in vivo with sufficient resolution. Because of its embryological origin, the retina is part of the central nervous system (CNS) and shares many anatomical, developmental, and functional features with the brain, making it a potential proxy for studying CNS disorders in living patients^1^. The retina is highly structured and organized into distinct layers containing cell bodies and neuropil^2^. It can be examined non-invasively with much higher resolution than the brain by using methods such as optical coherence tomography ^3^ and electroretinography.

Retinal investigations across major neuropsychiatric and neurological disorders have identified microstructural and electrophysiological alterations in schizophrenia (SCZ), bipolar disorder (BD), major depressive disorder (MDD),^4–11^ multiple sclerosis (MS),^12^ Alzheimer disease (AD),^13^ Parkinson disease (PD),^14^ and stroke^15–18^. For example, meta-analyses show reduced thicknesses of the retinal nerve fiber layer (RNFL) and the ganglion cell–inner plexiform layer (GCIPL)^4,19,20^ in individuals with SCZ compared with healthy controls. In MS, where RNFL and GCIPL thinning is also observed, retinal imaging is already used in clinical practice to support diagnosis and disease monitoring^12,21^.

However, the mechanistic underpinnings of retinal alterations in CNS disorders are poorly understood. Hypothetically, shared genetically driven mechanisms could directly affect retinal structures in brain diseases^1,7,22–29^ in addition to secondary mechanisms such as retrograde degeneration in MS^30^ and stroke^15^ or the effect of comorbidities in psychiatric disorders^31^. Previous studies have identified genetic overlap between retinal structures and CNS disease traits^25,29^ and have shown an association between polygenic disease risk and retinal alterations in SCZ,^7,26,32^ AD,^27^ and PD^27^. However, the cellular and molecular mechanisms that drive the link between the retina and CNS traits remain unknown.

The integration of summary statistics of genome-wide association studies (GWASs) and single cell transcriptomic data via gene set analysis tools such as MAGMA has been shown to be a powerful approach to elucidate pathophysiological mechanisms and identify relevant cell types in polygenic complex traits^33–35^. In this study, we mapped the polygenic risk of major neuropsychiatric and neurological disorders known to be associated with retinal alterations to retinal single cell gene expression data. Our aim was to identify which retinal cell types are associated with common variant genetic risk findings in major psychiatric and neurological disorders. Moreover, we explored which genetically driven pathophysiological mechanisms converge in retinal cells, and whether the high risk loading for SCZ found in amacrine cells (ACs) could be translated back to human retinal structure in a large population-based cohort.

## Methods

### Identification of cell type associations

By considering single cell and single nucleus retinal transcriptomic data sets from humans,^36–38^ non-human primates (NHPs),^39^ and mice,^40^ we calculated the specificity of each gene to different retinal cell types on the basis of the gene’s expression levels in those cell types (**eMethods** in **Supplement 1**). In addition, we assigned single nucleotide polymorphisms from GWAS summary statistics for stroke,^41^ AD,^42^ PD,^43^ MS,^44^ MDD,^45^ BD,^46^ and SCZ^47^ to nearby genes (**eMethods** in **Supplement 1**). Then, we performed a MAGMA cell type enrichment analysis^33,34^ (**Figure 1a**; **eMethods** in **Supplement 1**) to evaluate whether genetic risk is preferentially enriched in distinct cell types, as indicated by stronger genome-wide association signals in more cell type-specific genes.

**Figure 1:**
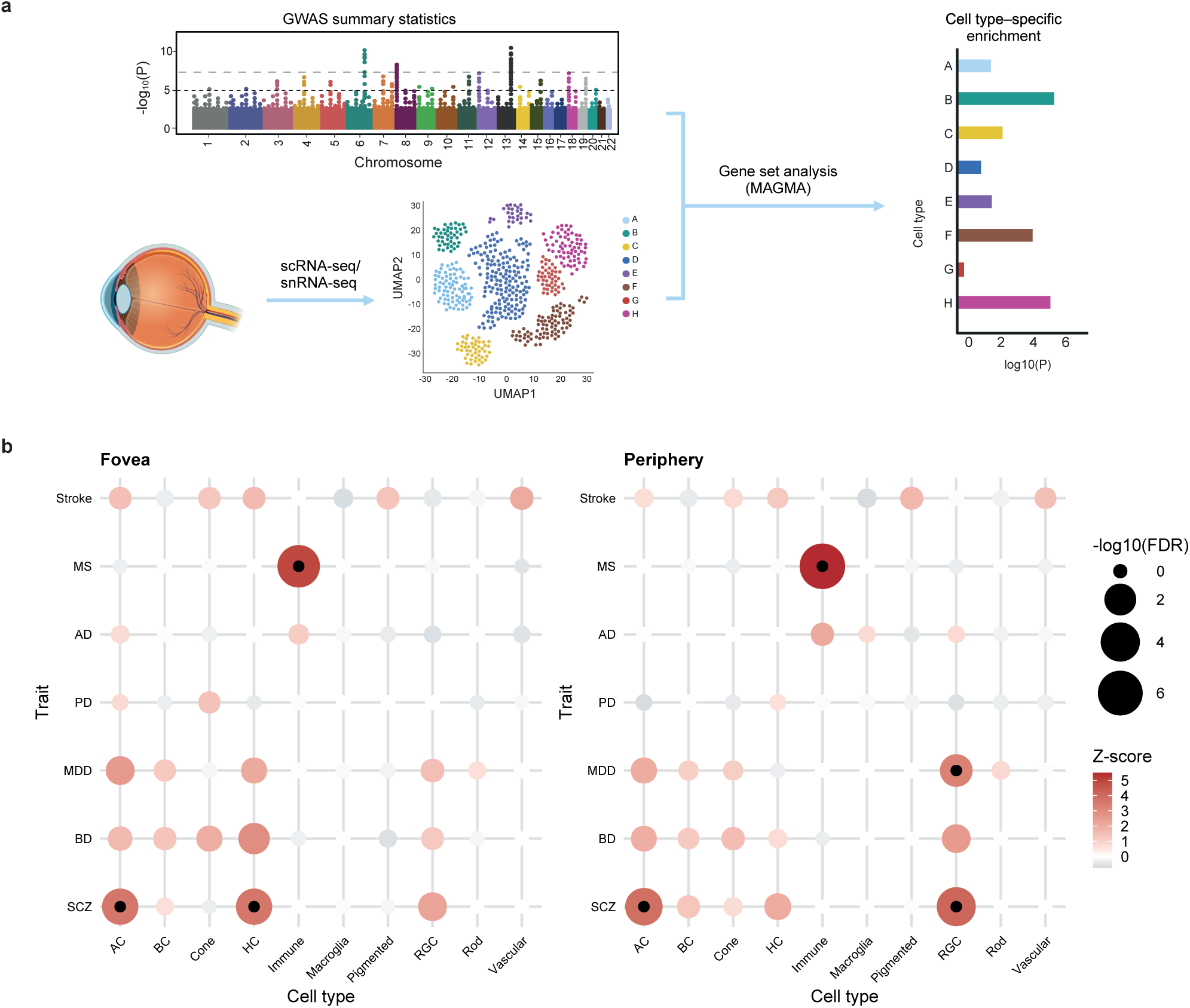
Gene set analysis reveals enrichment of genetic risk for major neurological and neuropsychiatric disorders in retinal cells. **a,** Schematic overview of the analytical approach. Single nucleotide polymorphisms from genome-wide association studies (GWASs) for several neurological and neuropsychiatric disorders were assigned to their nearby genes. In addition, specificity for each gene to retinal cell types was calculated from scRNA-seq expression data. Then, a MAGMA gene set analysis was performed to calculate the enrichment between genetic risk and the top 10% of genes that had the highest cell type specificity for each cell type. **b,** Dotplots illustrating the associations between retinal cell types and GWAS traits; dotplots were based on human scRNA-seq data^36^ for the fovea (left) and the peripheral retina (right). *P* values were adjusted for the false discovery rate within each trait. Inner black dots indicate statistical significance after additional Bonferroni correction for the seven investigated traits. The size of the outer dots represents the false discovery rate adjusted MAGMA *P* value. The color scale illustrates the MAGMA Z-score. *Abbreviations:* AC, amacrine cell; AD, Alzheimer disease; BC, bipolar cell; BD, bipolar disorder; FDR, false discovery rate-adjusted *P* value; GWAS, genome-wide association study; HC, horizontal cell; MDD, major depressive disorder; MG, Müller glia; MS; multiple sclerosis; PD, Parkinson disease; RGC, retinal ganglion cell; RPE, retinal pigment epithelium; scRNA-seq, single-cell RNA sequencing; SCZ, schizophrenia; snRNA-seq, single-nucleus RNA sequencing.

### Gene Ontology analyses

To gain insights into the neurobiological mechanisms through which polygenic risk acts in the revealed cell types we analyzed the enrichment of biologically relevant gene sets in the lists of genes that are associated with traits of interest and specifically expressed in trait-relevant cell types. To this end, we performed gene set enrichment analyses (GSEAs) with Enrichr^48–50^ and preranked GSEA^51^ (**eMethods** in **Supplement 1**).

### Validation and translation in a population-based cohort

We used population data from the UK Biobank (UKBB; www.ukbiobank.ac.uk) to validate and translate our top cellular findings. The population data included Caucasian British and Irish individuals without a diagnosis of SCZ or schizotypal or delusional disorder (**eFigure 7** in **Supplement 1**). We computed a polygenic risk score (PRS) for SCZ (SCZ-PRS) from the latest release of genome-wide association data^47^. By using robust regression, we tested whether genetic susceptibility to SCZ is associated with alterations of the RNFL, GCIPL, and inner nuclear layer (INL) thickness (**eMethods** in **Supplement 1**).

## Results

### Cell type-specific enrichment of GWAS traits in retinal cells

Our analyses were based on the assumption that the polygenic risk burden for a complex brain disorder accumulates not only in the brain cell types that are mainly relevant for the clinical phenotype^34,35^ but also in tissues and cell types affected by the same genetic architecture and, presumably, by the same mechanisms that are relevant for the said clinical phenotype. A disease-relevant cell type will show an accumulation of expressed genes with high genetic risk loading, and if the functions of said cell type are relevant for pathobiology, the signal will increase with the specificity of expression. Cell types in other tissues and contexts that express the same genes may be affected by the same genetic risk and produce a different or similar pathophysiological state.

To investigate such associations of major polygenic brain disorders in retinal cell types, we performed a MAGMA cell type enrichment analysis by integrating GWAS summary statistics with single-cell RNA sequencing (scRNA-seq) data from light-responsive human post mortem retinas^36^ (**Figure 1b**; **eTable 1** and **2** in **Supplement 2**). The analysis revealed that the expression profile of ACs was significantly enriched for genetic risk of SCZ in both the fovea and the peripheral retina. Less consistently, SCZ also mapped to other retinal neurons in humans, i.e., to horizontal cells (HCs), but only in the foveal region, and to retinal ganglion cells (RGCs), but only in the peripheral retina. After controlling the expression specificity of each gene for ACs in MAGMA, the associations with HCs and RGCs with SCZ were still significant. This finding indicates that the observed inconsistent enrichment of HCs and RGCs cannot be solely explained by genetic overlap between retinal neurons (**eTable 3** and **4** in **Supplement 2**).

In MDD, we found a link to peripheral retinal ganglion cells (RGCs), albeit with a weaker effect than for SCZ, and in MS, we identified a significant enrichment with foveal and peripheral retinal immune cell populations. In stroke, AD, PD, and BD, analyses showed no significant cell type associations with genetic risk.

To exclude confounding by data set-specific batch effects, we validated our findings in an independent human single-nucleus RNA sequencing (snRNA-seq) data set^38^. This analysis confirmed significant enrichment for SCZ risk in retinal ACs and for MS risk in retinal microglia (**eFigure 1a** in **Supplement 1**; **eTable 5** in **Supplement 2**).

ACs represent a highly diverse population of interneurons that, with a few exceptions,^52^ release inhibitory neurotransmitters^53^. To investigate whether the enrichment between SCZ risk and ACs was driven by a particular subtype, we performed additional cell type enrichment analyses within the comprehensively annotated AC snRNA-seq subset from the Human Retinal Cell Atlas^38^. This analysis showed no particular association with a specific subtype; this was the case when we considered the major AC subpopulations classified by the inhibitory neurotransmitters they express (GABAergic, glycinergic, both GABAergic and glycinergic, or non-GABAergic non-glycinergic; **eFigure 1b** in **Supplement 1**; **eTable 6** in **Supplement 2**) and when we considered the detailed resolution of all 73 AC subtypes (**eFigure 1c** in **Supplement 1**; **eTable 7** in **Supplement 2**).

### Expression of SCZ risk genes in the developing human retina

Previous studies implicate prenatal neurodevelopment in SCZ,^54,55^ so we explored whether the enrichment of GWAS traits exhibits different patterns in snRNA-seq data from fetal retinal cells^37^. In this evaluation, we found a robust enrichment for SCZ risk in fetal ACs (**eFigure 2** in **Supplement 1**; **eTable 8** in **Supplement 2**). Fetal retinal cell types showed no association with any other disorder.

**Figure 2:**
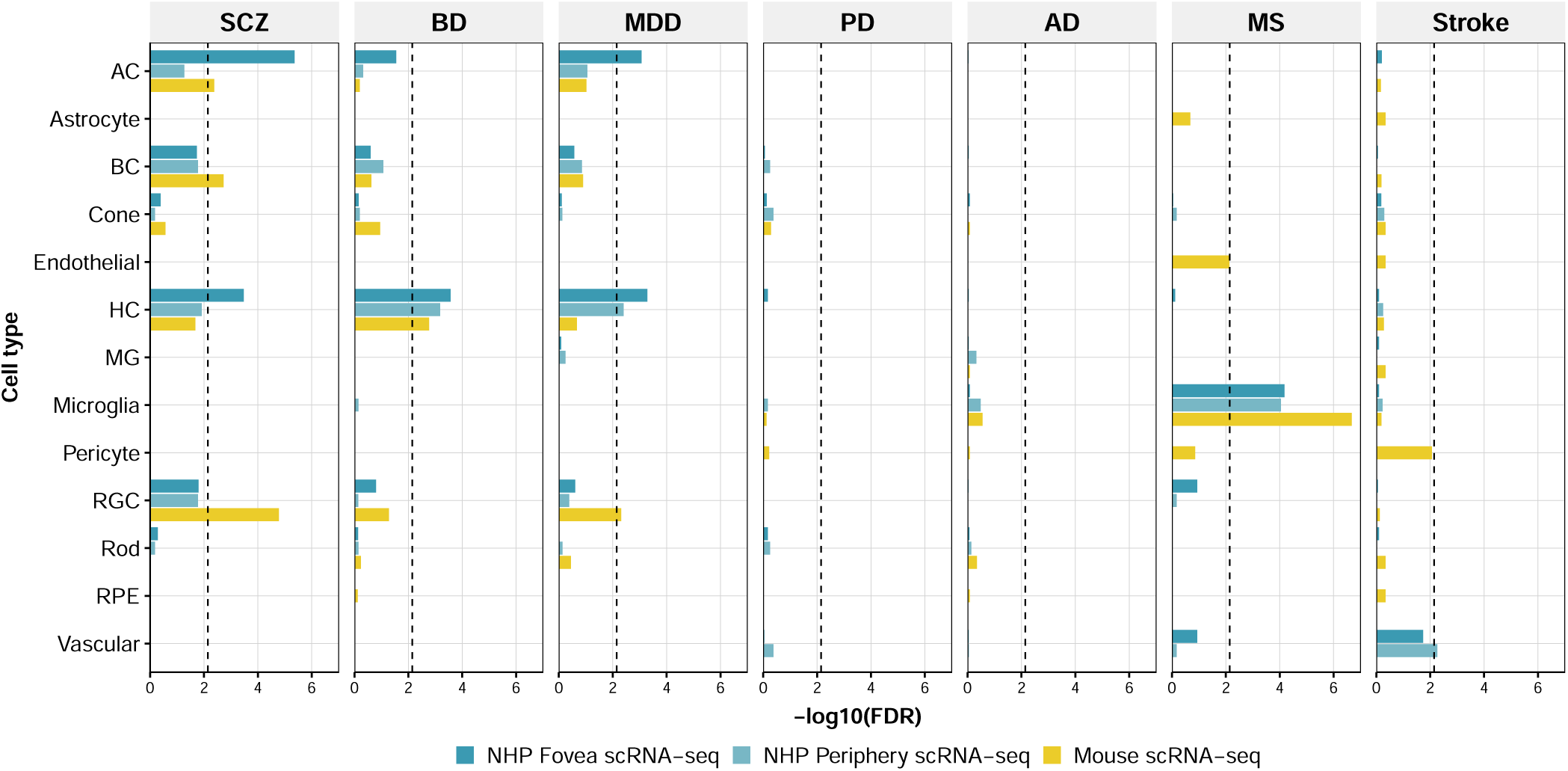
Associations between traits and retinal cell types are robust across mammalian species. Bar plots illustrating the log-transformed MAGMA *P* values (false discovery rate-corrected across each trait) for the association between retinal cells and schizophrenia, bipolar disorder, major depressive disorder, Parkinson disease, Alzheimer disease, multiple sclerosis, and stroke in the non-human primate *macaca fascicularis*^39^ and in mice^40^. The dashed lines represent the Bonferroni-corrected significance level of 0.05/7. Note that vascular cells, retinal pigment epithelium, pericytes, endothelial cells, and astrocytes were not similarly annotated in the two species. *Abbreviations:* AC, amacrine cell; AD, Alzheimer disease; BC, bipolar cell; BD, bipolar disorder; FDR, false discovery rate-adjusted *P* value; HC, horizontal cell; MDD, major depressive disorder; MG, Müller glia; MS; multiple sclerosis; NHP, non-human primate; PD, Parkinson disease; RGC, retinal ganglion cell; RPE, retinal pigment epithelium; SCZ, schizophrenia.

### Validation of retinal enrichment across species

To validate the associations identified in human transcriptomes and look for conservation of these findings in mammalian model organisms, we repeated the cell type association analysis with scRNA-seq data from non-human primates (NHPs)^39^ and mice^40^ (**Figure 2**; **eTable 9**–**11** in **Supplement 2**). These analyses confirmed the robust association of SCZ with ACs and revealed significant enrichment in foveal ACs of NHPs and in mice ACs, but not in peripheral ACs of NHPs. Moreover, we found inconsistent enrichment of the SCZ trait in the transcriptome of other retinal neurons of both species, namely bipolar cells (BCs) and RGCs of mice and foveal HCs of NHPs. For BD, the results showed a significant enrichment in HCs in both species. MDD was inconsistently enriched in ACs, HCs, and RGCs. In line with the findings in the human retina, genetic risk for MS was enriched in microglia from both species. Notably, in stroke the enrichment of the genome-wide association signal in peripheral retinal vascular cells also reached statistical significance in NHPs.

In summary, in analyses of transcriptomic data from different species and developmental stages we found that SCZ was robustly enriched in retinal ACs and that MS robustly mapped to immune cell populations but not to neural cells.

### Enriched genes converge on distinct gene sets

To reveal the biological mechanisms behind the identified trait–cell type associations, we used Enrichr^48–50^ to perform GSEAs with those cell type-specific genes that were significantly associated in the human transcriptome with the trait of interest (**Figure 3**; **eFigure 3** and **4** in **Supplement 1**).

**Figure 3:**
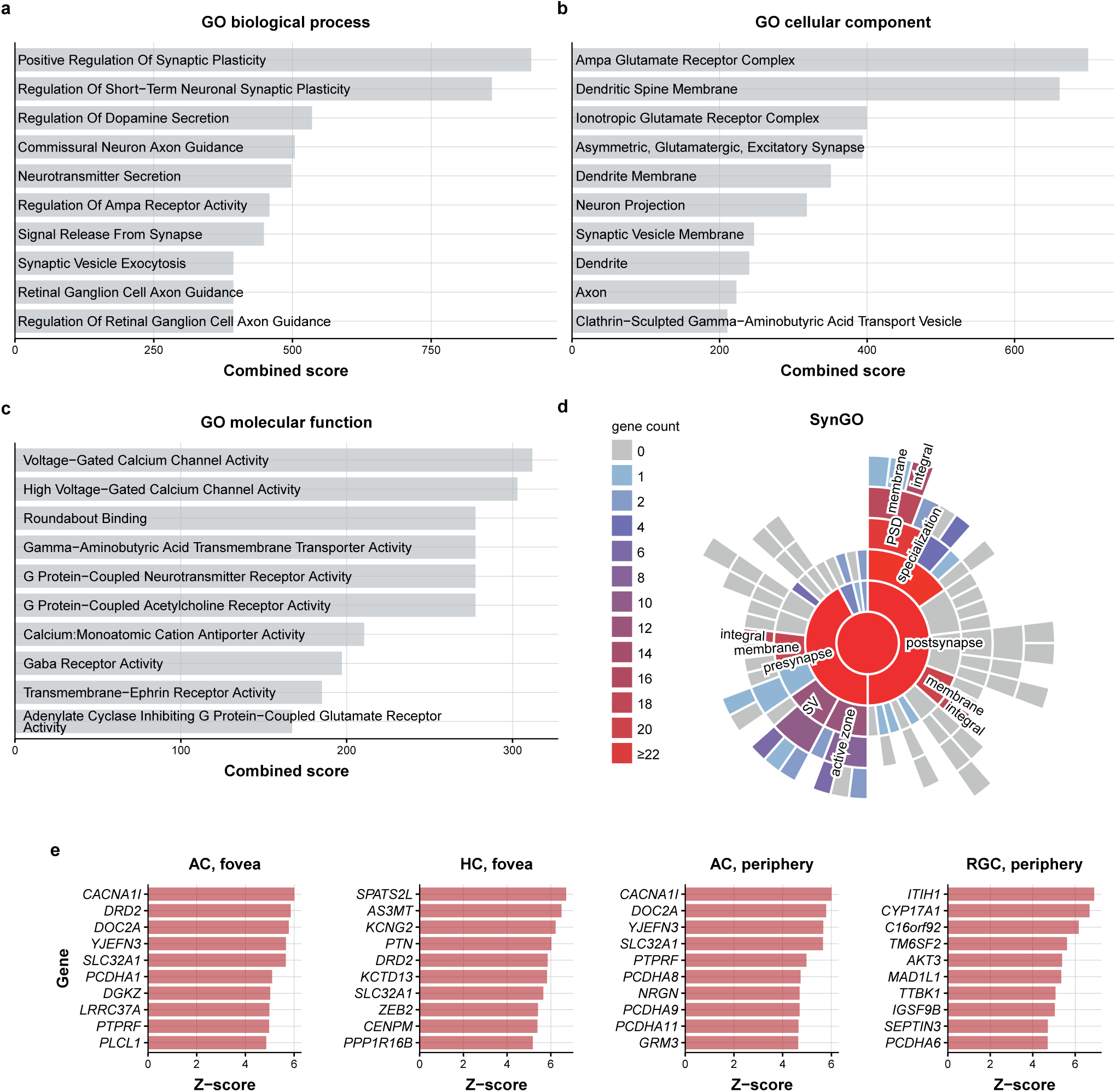
Enrichment analysis implicates disrupted retinal synapses in schizophrenia. **a–c,** Bar plots displaying the top 10 gene sets from Gene Ontology (GO) biological process (**a**), GO cellular component (**b**), and GO molecular function (**c**) that were overrepresented among those genes that were specifically expressed in amacrine cells and significantly associated with schizophrenia (SCZ), as obtained with Enrichr. Specifically, the enrichment analysis was based on the overlap of 226 genes that were in the top specificity decile for both foveal and peripheral amacrine cells and had an unadjusted MAGMA gene-level *P* value of less than 0.05. For additional analyses for each relevant cell type and region separately, see **e**Figure 4. **d,** Sunburst plot from SynGO based on 84/226 genes that were significantly associated with SCZ on the gene level, had a cellular component annotation in SynGO, and were specifically expressed in both foveal and peripheral amacrine cells in the Cowan et al. data set^36^. Colors indicate gene count per term, including child terms. Fields with 10 or more genes mapping to them or their child terms are annotated. **e–h,** Bar plots illustrating the 10 genes with the highest MAGMA Z-scores for SCZ within the top 10% cell type-specific genes of each cell type that was significantly associated with genetic risk for SCZ in the human data set of Cowan et al.^36^ *Abbreviations:* AC, amacrine cell; HC, horizontal cell; RGC, retinal ganglion cell.

**Figure 4:**
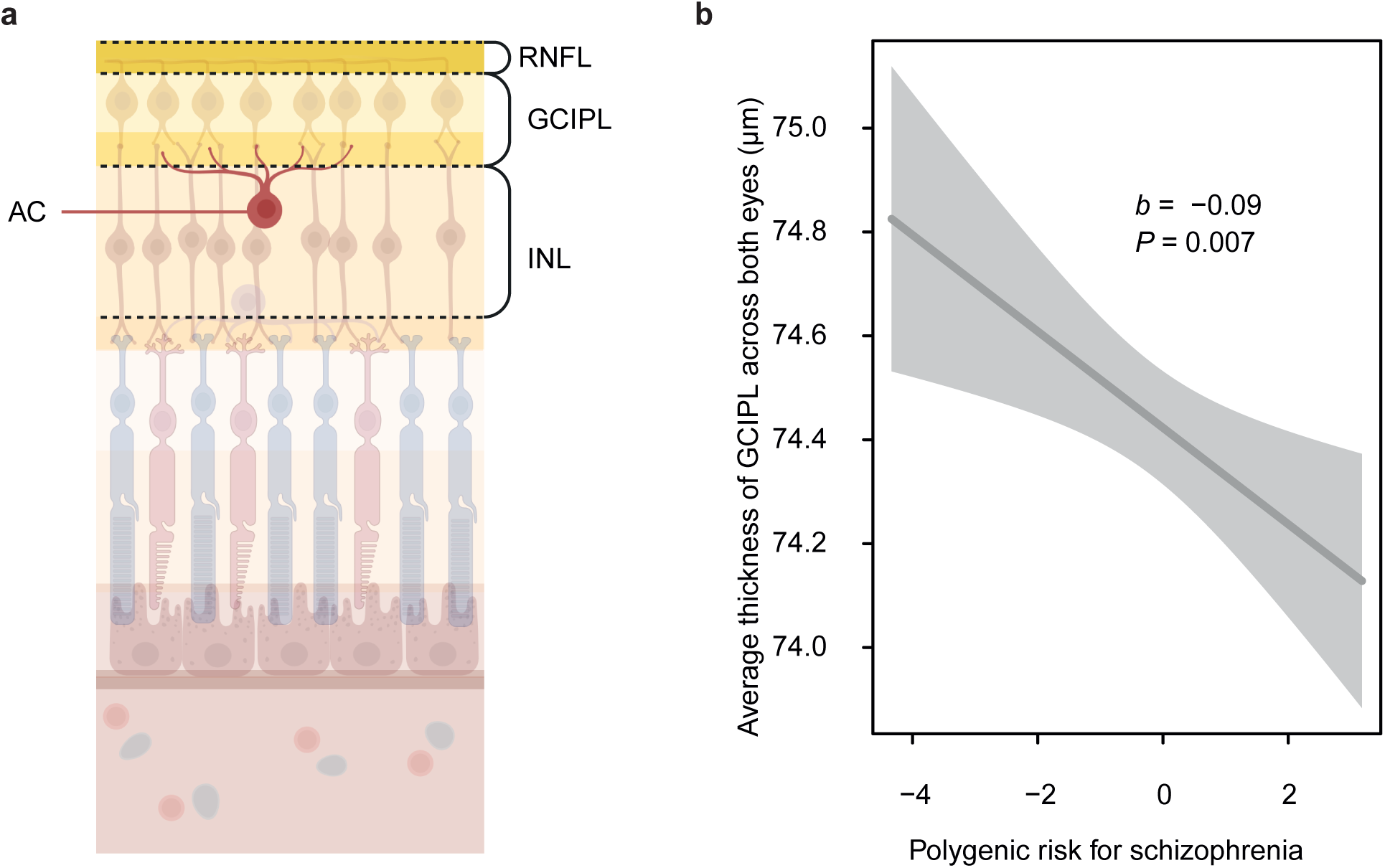
Ganglion cell–inner plexiform layer thickness is associated with polygenic risk for schizophrenia in a population-based cohort. **a,** Schematic illustration of the retinal layers (retinal nerve fiber layer, ganglion cell–inner plexiform layer [GCIPL], inner nuclear layer) that were investigated within the population-based cohort and of the anatomical location of the amacrine cells. **b,** Partial robust regression plot between GCIPL thickness (averaged across both eyes) and polygenic risk scores for schizophrenia in the UK Biobank cohort (*n* = 36,349). Solid line represents the regression line, and the complementary shaded area corresponds to the 95% CI. *Abbreviations:* AC, amacrine cell; GCIPL, ganglion cell–inner plexiform layer; INL, inner nuclear layer; RNFL, retinal nerve fiber layer.

In SCZ, we found broad involvement of gene sets related to synapse biology, including gene sets involved in synaptic plasticity and in calcium and glutamatergic signaling in ACs (**Figure 3a**–**c**). According to SynGO^56^ annotation, the SCZ risk genes with high specificity for ACs were relevant for both pre and postsynaptic structures, such as the presynaptic membrane and postsynaptic density membrane, respectively (**Figure 3d**). To validate our findings with another method, we performed a preranked GSEA^51^; the results of this analysis also pointed to neuronal and synaptic function, and in addition, to cell adhesion (**eFigure 3**).

Among the genes specifically expressed in ACs, HCs, and RGCs with the strongest association to SCZ were *CACNA1I,* which encodes the calcium voltage-gated channel subunit alpha1 I; *DOC2A*, which is relevant for Ca^2+^-dependent neurotransmitter release^57^; the dopamine receptor D2 (*DRD2*), which is the common target of antipsychotics^58^; and *SLC32A1*, which codes for a protein involved in GABA and glycine uptake into synaptic vesicles^59^ (**Figure 3e**; **eTable 12–15** in **Supplement 3**).

In MDD, the GSEAs yielded gene sets involved in neuronal biology, but with less involvement in synapse biology (**eFigure 5**; **eTable 16** in **Supplement 3**). In MS, the enrichment in retinal immune cells converged onto immunological gene sets, especially those involved in cell-mediated immunity (**eFigure 6** in **Supplement 1**), with interleukin 2 receptor subunit alpha (*IL2RA*) as the most strongly associated gene (**eTable 17** and **18** in **Supplement 3**).

### Schizophrenia polygenic risk and retinal imaging

To translate the most consistent genetic association of the SCZ trait with ACs to the human retinal microstructure in vivo, we explored the association between SCZ-PRSs and retinal layer thicknesses in 36,349 individuals of the UKBB cohort (**eFigure 7** in **Supplement 1**; **Figure 4**). Robust linear regression revealed a significant association of higher SCZ-PRS with reduced GCIPL thickness (*b* = −0.09; 95% CI [−0.16, −0.03]; *P* = 0.007; **Figure 4b**) but no significant associations with RNFL (*P* = 0.481) and INL (*P* = 0.865) thickness (**eTable 19** in **Supplement 4**).

## Discussion

By integrating GWAS summary statistics and single cell transcriptomics, we showed that the polygenic risk of different CNS disorders converges on distinct cell types and biological functions in the retina. Most notably, we found robust associations between SCZ risk and ACs and between MS risk and retinal immune cells, which were conserved across different mammalian species. Interestingly, the enrichment of SCZ in ACs was also detectable in fetal retinal tissue, underlining that the retina reflects the neurodevelopmental component in SCZ pathogenesis^54^.

Less consistently, we detected associations of HCs, RGCs, and BCs with SCZ in several human retinal data sets from different anatomical regions and in transcriptomic data from mice and NHPs. Some inconsistencies between species might be explained by the fact that although most cell types are conserved, marked anatomical differences also exist, not least the absence of a fovea in mice^2,60^. Moreover, technical variations in tissue preparation and scRNA-seq processing may also have contributed to these inconsistencies.

ACs are mostly GABAergic or glycinergic interneurons with diverse functions in visual signal processing^53^. They provide feedback on BCs, whose response to light has been found to be reduced in SCZ,^7,9^ and synapse with RGCs and with each other within the inner plexiform layer (IPL)^53,61,62^. Previous brain-based post-GWAS analyses highlighted a disturbed synapse biology in SCZ that also affects GABAergic interneurons^47,63^. Here, we found that the polygenic disease risk also converges on gene sets that are highly relevant for synaptic function in ACs. In particular, the functions in which these gene sets are involved include synaptic plasticity and neurotransmitter biology, which have been identified to be dysregulated in SCZ^64,65^.

We translated these genetically driven cellular findings to the microstructural level by demonstrating an association of higher SCZ-PRS with GCIPL thinning in a non-schizophrenic population-based cohort, the UKBB. This finding, which was based on the latest SCZ GWAS^47^, aligns with a very recent analysis of UKBB data^32^ using a different GWAS discovery sample^66^. The GCIPL comprises the ganglion cell layer, which contains RGC bodies, and the complex neuropil of the IPL^67^. Given that our study found no association for the RNFL, the axonal layer of RGCs, it is less likely that the revealed association of GCIPL thickness with SCZ-PRS is driven by alterations of RGCs. Instead, it might reflect synaptic impairments within the IPL in SCZ. The somas of ACs, BCs, and HCs in the INL appeared not to be affected. This finding aligns with a recent study from our group which highlighted pronounced thinning of the IPL^7^ in SCZ spectrum disorders. The present study indicates that synapse biology in SCZ, which is disturbed in the brain^46^, is also altered in the retina and thereby might contribute to the microstructural alterations in the complex neuropil of the IPL^67^ in patients with SCZ. Given the difficulties in investigating synaptic pathology in the brain itself in vivo in humans,^63,68^ our findings highlight the retina as a unique opportunity to directly image microstructures that could reflect synaptic pathology in the CNS.

In addition, our study yielded inconsistent evidence of an association between several types of retinal neurons and affective disorders. Considering the robust association of retinal neurons with genetic SCZ risk, the less consistent enrichment of BD and MDD in retinal cells might reflect their genetic overlap with SCZ^69^ within the spectrum of neuropsychiatric disorders^70^.

With respect to neurological disorders, only MS risk was robustly enriched in a distinct cell population. The link between MS and retinal immune cells and microglia is consistent with the known enrichment of MS risk genes in brain microglia^44^ and could suggest a role of resident immune cells in retinal thinning in MS^71^. In addition, a recent study demonstrated genetic pleiotropy of retinal structures with AD and PD^29^. An increased AD-PRS has been linked to retinal changes such as a thicker IPL and INL, and an increased PD-PRS, to a thinner outer plexiform layer^27^. However, other studies failed to find relevant genetic associations between AD and the thicknesses of the RNFL or GCIPL^72^ or between PRSs for retinal layer thicknesses and incident or prevalent AD or PD^73^. On a cellular level, postmortem findings point to a reduction of dopaminergic ACs in PD^74^. Conversely, AD, which is genetically associated with brain microglia,^42,75^ has been linked to dysfunction of retinal glia^76–79^. Despite these previously described associations of retinal cell populations with AD and PD, our study found no significant enrichment of AD or PD with specific cell types. Of note, the negative finding regarding a genetic link between retinal microglia and AD is consistent with previous work^22^.

In summary, among several neurological and psychiatric diseases that were previously linked to retinal alterations, only the polygenic architecture of SCZ and MS robustly mapped to specific and distinct retinal cell populations, indicating distinct disease- and cell type-specific mechanisms underlying the reported retinal alterations in SCZ and MS.

### Limitations

Our approach has several limitations. Because we focused on highly cell type-specific genes, effects of genes with broader, tissue-wide, or pan-neuronal expression patterns are underrepresented. Nevertheless, this approach enables better interpretability because by minimizing overlap between cell types, it yields fewer significant results^80^. Second, technical limitations of scRNA-seq and snRNA-seq may lead to underrepresentation of transcripts exported to distal cellular compartments^34,81^. Moreover, expression patterns may change with age^35^. The donor age range of the Cowan et al. retina atlas was between 50 and 80 years,^36^ which is far beyond the typical age of onset of SCZ^82^. However, we applied our analyses to retinas from two developmental stages in humans (adults and fetuses), and the finding for ACs in SCZ was robust across both these developmental stages and across transcriptomic data sets from different species.

## Conclusions

This study demonstrates that the polygenic risk for SCZ and MS converges on specific cell populations within the retina, and it provides evidence that to a certain degree, the retina mirrors genetically driven pathophysiological mechanisms in SCZ and MS. The distinctly layered microstructure of the retina could serve as a window to the brain and enable the evaluation of cellular and synaptic pathology in vivo in affected patients, e.g., by assessing structural changes of the GCIPL in SCZ. In sum, this work highlights the retina’s potential to serve as an accessible proxy for studying the neurobiology of CNS disorders.

## Data availability

Summary statistics for AD, BD, MDD, and SCZ were obtained from https://pgc.unc.edu/for-researchers/download-results/. Summary statistics for PD and stroke were downloaded from the NHGRI-EBI catalogue under accession codes GCST90275127 and GCST90104534 (https://www.ebi.ac.uk/gwas/studies/). MS summary statistics were sourced from the Data Access Committee of the International Multiple Sclerosis Genetics Consortium (IMSGC) via https://imsgc.net/?page_id=31. Chan Zuckerberg CELLxGENE Discover was used to download retinal scRNA-seq data of Cowan et al. (https://cellxgene.cziscience.com/collections/2f4c738f-e2f3-4553-9db2-0582a38ea4dc), the MRCA (https://cellxgene.cziscience.com/collections/a0c84e3f-a5ca-4481-b3a5-ccfda0a81ecc), and the HRCA (https://cellxgene.cziscience.com/collections/4c6eaf5c-6d57-4c76-b1e9-60df8c655f1e). The NHP expression data of Peng et al. were downloaded from the Single Cell Portal (https://singlecell.broadinstitute.org/single_cell/study/SCP212/). HomoloGene is available at https://www.ncbi.nlm.nih.gov/datasets/gene/. g:Profiler is available at https://biit.cs.ut.ee/gprofiler/gost). MSigDB is available at https://www.gsea-msigdb.org/gsea/msigdb/. Information on how to access data from the UK Biobank is available at https://www.ukbiobank.ac.uk/enable-your-research/apply-for-access.

## Code availability

SynGO is available at https://www.syngoportal.org/. Enrichr is available at https://maayanlab.cloud/Enrichr/. The MAGMA.Celltyping R package is available at https://github.com/neurogenomics/MAGMA_Celltyping.

## Acknowledgments

The authors thank Jacquie Klesing, BMedSci (Hons), Board-certified Editor in the Life Sciences (ELS), for editing assistance with the manuscript. Figure 1a and Figure 4a were created with Biorender.com.

## Author contributions

Dr Rabe had full access to the UK Biobank data used in the study. Mr Boudriot had full access to all other data in the study except the UK Biobank data. Both authors take responsibility for the integrity of the respective data and the accuracy of the data analysis.

*Concept and design:* Boudriot, Raabe, Stephan.

*Acquisition, analysis, or interpretation of data:* Boudriot, Papiol, Raabe, Rabe, Smigielski, Stephan.

*Drafting of the manuscript:* Boudriot, Papiol, Raabe, Rabe, Smigielski.

*Critical revision of the manuscript for important intellectual content:* All authors.

*Statistical analysis:* Boudriot, Papiol, Raabe, Rabe, Smigielski, Stephan.

*Obtained funding:* Boudriot, Falkai, Homan, Raabe, Rossner, Schmitt.

*Administrative, technical, or material support:* Falkai, Homan.

*Supervision:* Raabe.

## Conflict of Interest Disclosures

Prof Rossner is shareholder of, and Dr Stephan is employed by Systasy Bioscience GmbH, Munich, Germany. Prof Falkai received speaking fees from Boehringer-Ingelheim, Janssen, Otsuka, Lundbeck, Recordati, and Richter and was a member of the advisory boards of these companies and Rovi. Prof Homan has received grants and honoraria from Novartis, Lundbeck, Mepha, Janssen, Boehringer Ingelheim, and Neurolite outside of this work. No other disclosures were reported.

## Funding/Support

Mr Boudriot received funding from the Pesl Alzheimer Foundation (Pesl-Alzheimer-Stiftung; 2024– 2025) and the Foundations of the Medical Faculty of the LMU Munich (Stiftungen zugunsten der Medizinischen Fakultät – Cluster 2; 2024–2025). Prof Falkai was supported by the Federal Ministry of Education and Research (Bundesministerium für Bildung und Forschung [BMBF]) within the initial phase of the German Center for Mental Health (DZPG; grant FKZ 01EE2303A and 01EE2303F) and within the Era-Net Neuron project GDNF_UpReg (FKZ 01EW2206). Prof Homan is supported by a NARSAD grant from the Brain & Behavior Research Foundation (28445), by a Research Grant from the Novartis Foundation (20A058), and by an ERC Synergy Grant (101118756). Dr Papiol was supported by European Union’s Horizon Europe research and innovation program under grant agreement No 101057454 (Psych-STRATA). Dr Raabe was supported by the Munich Clinician Scientist Program (MCSP) of the Medical Faculty, LMU Munich (FöFoLe 009/2019 and Advanced Track 01/2021), the Lisa Oehler Foundation (Dr. Lisa Oehler-Stiftung; 2022–2024), and the Pesl Alzheimer Foundation (Pesl-Alzheimer-Stiftung; 2024–2025).

## Role of the Funder/Sponsor

The funding sources had no role in the design and conduct of the study; collection, management, analysis, and interpretation of the data; preparation, review, or approval of the manuscript; and decision to submit the manuscript for publication.

## Supplementary information

Supplement 1: eMethods and eFigure 1–7.

Supplement 2: eTable 1–11.

Supplement 3: eTable 12–18.

Supplement 4: eTable 19.

