## Supplement 1 for "Genetic analysis of retinal cell types reveals synaptic pathology in schizophrenia"

***Boudriot* et al.**

Supplement 1 (this document): eMethods and eFigure 1–7.

Supplement 2: eTable 1–11.

Supplement 3: eTable 12–18.

Supplement 4: eTable 19.

### eMethods

#### Genome-wide association study summary statistics and gene mapping

Genome-wide association study (GWAS) summary statistics for stroke[1], Alzheimer disease (AD)[2], Parkinson disease (PD)[3], multiple sclerosis (MS)[4], major depressive disorder (MDD)[5], bipolar disorder (BD)[6], and schizophrenia (SCZ)[7] were standardized for subsequent analyses by using the *format_sumstats* function of the *MungeSumstats*[8] package in *R*[9]. SNPs were assigned to nearby genes on basis of their physical location by using the function *map_snps_to_genes* from package *MAGMA.Celltyping*[10,11] and considering SNPs within 35 kb upstream and 10 kb downstream of each gene.

#### Retinal transcriptome

As a reference for gene expression in the foveal and peripheral regions of the human retina, we used the human retina scRNA-seq data set provided by Cowan et al.[12] This data resource was selected because of its high quality and methodological rigor. The authors reduced post-mortem ischemic times to less than five minutes, and additional electrophysiological investigations ensured light responsiveness of post-mortem tissue[12]. For replication, we used the large-scale snRNA-seq data set of the Human Retinal Cell Atlas (HRCA), version 1[13]. Notably, for analyses performed in the complete data set of all cell types, the HRCA was randomly downsampled to a maximum 5000 cells per major class to facilitate computations. Furthermore, we used snRNA-seq data of retinas from human fetuses that were 9 to 24 weeks post fertilization[14]. Classification of cells into major retinal cell classes was based on annotations provided by the respective authors.

To assess the generalizability of our findings across mammalian species, we also performed analyses in scRNA-seq data of the foveal and peripheral retina of the non-human primate (NHP) *macaca fascicularis*[15] and in retinal scRNA-seq data of wild-type mice; for the latter, we used the integrated Mouse Retina Cell Atlas (MRCA)[16]. Mouse genes were converted to their human orthologs by referring to HomoloGene[17], and macaque genes, by referring to g:Profiler[18].

For each data set, we calculated the cell type expression specificity of each gene to different retinal cell types by dividing the mean expression in the respective cell type by the mean expression in all cell types[10]. This was done with the function *generate_celltype_data* of the package *EWCE*[19].

#### Identification of disease-relevant cell types

To identify retinal cell types with expression patterns significantly enriched for GWAS risk loci associated with stroke, AD, PD, MS, MDD, BD, and SCZ, we used the gene set analysis method MAGMA v1.10 performed with the *MAGMA.Celltyping R* package (v2.0.11). The method is based on the assumption that if a cell type is involved in the trait of interest, the genes most specifically expressed in that cell type will also have a stronger association with the trait. This specific use case of MAGMA was developed and is described in detail by Skene et al.[10]. A general description of MAGMA can be found in [11]. MAGMA was chosen because it is generally considered to be more powerful than other established methods[20].

In short, MAGMA was used to calculate a cumulative *P* value for each expressed gene as a metric for its association with the trait of interest (gene level *P* value). As described elsewhere, this computation is based on the SNPs assigned to the respective genes and their individual *P* values in the GWAS of interest[11]. This step was integrated into the *map_snps_to_genes* function from the *MAGMA.Celltyping* package[10,11]. The gene-level *P* value was then converted to a Z-score, which measures the strength of the association between each gene and the trait[11]. In a second step, a competitive gene set analysis was computed to test whether the 10% most cell type-specific genes were more strongly associated with the trait than the rest of the genes were. This was done with the "Top 10%" mode of the *celltype_associations_pipeline* function from the *MAGMA.Celltyping* package[10,11]. Note that this method disregards any differences in association strength within the top 10% gene set and any meaningful association in the remaining nine deciles of the expressed genes in the cell type in question.

Cell type-level *P* values were adjusted for the false discovery rate (FDR)[21] within each GWAS trait. The overall significance level was additionally adjusted for the total number of traits by using Bonferroni correction ($\alpha = 0.05/7$). This was done separately for each transcriptomic data set.

To eliminate the possibility of confounding because of genetic overlap between cell types, we conducted an additional conditional analysis that could take gene-level cell type specificity in other cell types into account. This analysis was performed with the *calculate_conditional_celltype_associations* function from the *MAGMA.Celltyping* package. Since this analysis was only performed for SCZ, an FDR corrected *P* value of less than 0.05 was considered statistically significant.

#### Gene Ontology enrichment analyses

After identifying disease-relevant retinal cell types, we aimed to gain deeper insights into underlying biological mechanisms by performing additional Gene Ontology (GO)[22,23] enrichment analyses. To ensure robust results, we used multiple methods for these analyses. First, we used Enrichr[24-26] to examine the convergence of those genes within the top decile of cell-type specificity that were significantly linked to the trait of interest (as indicated by a MAGMA gene-level *P* value of less than 0.05) and GO gene sets. The Enrichr "combined score," which is calculated by multiplying the odds ratio by the negative natural logarithm of the *P* value derived from Fisher's exact test[26], was used as a ranking metric. The Enrichr analysis was performed separately for each cell type-GWAS pair with a statistically significant result in the gene set analysis of the human data set from Cowan et al.[12] For SCZ, we performed this analysis also only with genes that were in the top specificity decile in both regions (i.e., fovea and periphery) to minimize the possibility of spurious findings. To gain further insights into synaptic locations of the genes, the latter gene list was also visualized with SynGO[27] (**Figure 3d**).

As a second method, we performed GSEA with the *fgsea*[28] package in *R*, which applied an adaptive multi-level splitting Monte Carlo approach[29]. This analysis was also done separately for each cell type that was significantly associated with the respective trait. All genes within the top specificity decile and all GO terms with a gene set size of 15 to 500 genes were used as input. GO gene sets were obtained from MSigDb[30-32]. The MAGMA Z-score[11] was used as a ranking metric within the top specificity decile of genes. To retain only independent gene sets and ease interpretability, we applied the *collapsePathways* function from *fgsea*.

#### Retinal imaging

To investigate whether our most consistent finding of an enrichment of the SCZ trait in amacrine cells could be translated to the microstructural level, we used data from the UK Biobank (application 102266). We focused here on SCZ and not on MS, because in MS, the cell type enrichment analysis showed no enrichment in retina-specific cell types, only in immune cells and microglia within the retina. Retinal optical coherence tomography data of 65,484 participants, acquired between 2009 and 2010, were obtained from the Biobank. Macular volume scans were performed with a 3D OCT-1000 Mark II (Topcon, Japan) and segmented with the Topcon Advanced Boundary Segmentation (TABS) algorithm. We focused on the two retinal layers that are microstructurally linked to amacrine cells: the inner nuclear layer (INL; field IDs 28502, 28503), where the somas of ACs are located, and the ganglion cell–inner plexiform layer (GCIPL; field IDs 28504, 28505), which comprises the neuropil of BCs, RGCs, and ACs and RGC somas. Assuming that any changes in the GCIPL due to alterations in RGCs would also affect the retinal nerve finer layer (RNFL), which contains the RGC axons that exit the retina via the optic nerve, we examined also the RNFL (field IDs 28500, 28501). The complete image acquisition and segmentation procedure is described in detail elsewhere[33]. Images falling within the lowest 20% of image quality were excluded from the analysis.

#### Genetic data from the UK Biobank, and PRS calculation

Detailed descriptions of the genotyping and imputation procedures for the UK Biobank data can be found in the release documentation[34]. In summary, 487,409 blood samples were analyzed with two customized tagSNP arrays (the Applied Biosystems UK BiLEVE Axiom Array and the Applied Biosystems UK Biobank Axiom Array [Affymetrix, Santa Clara, CA, USA]), which share 95% of their markers. The data were imputed to the UK10K and 1000 Genomes Project Phase 3 reference panels: SHAPEIT3[35] was used for phasing, and IMPUTE2[36] for imputation. Additional data handling and quality control (QC) steps were performed according to a published processing pipeline[37] . To address population stratification, 10 genetic principal components were retrieved from the UK Biobank. Specifically, after SNP extraction and alignment, conversion from bgen to PLINK format, and removal of ambiguous SNPs (A/T, C/G; allele frequencies between 0.4 and 0.6), the data underwent SNP-level

QC (minor allele frequency < 0.005 and INFO score < 0.4) and sample-level QC (retaining individuals with a missing rate in autosomes ≤ 0.02, who were not outliers for genotype missingness or heterozygosity, not genetically related up to third-degree relatives, not sex-discordant, and of Caucasian British or Irish ethnicity according to genetic grouping). Individuals with ICD-10 diagnoses F20 to F29; those medicated with antipsychotics; those with diabetes related eye disorders, glaucoma, macular degeneration, or other serious eye conditions; and those with missing data in the variables of interest or covariates were excluded. Ultimately, a total of $n$ = 36,349 individuals with matched imaging-genetic data were included in the final analysis (see **eFigure 7** in **Supplement 1** for details). As a last step, polygenic risk scores for SCZ (SCZ-PRS) were computed for each individual by summing the risk alleles weighted by their estimated effect sizes[38] with the --score function in PLINK 2.0[39]. A linkage disequilibrium pruning was performed with a 200 kb genomic distance, $r^2 \geq 0.3$, and a 1-SNP (default) step size.

#### Robust regression between retinal layer thickness measures and SCZ-PRS

We used linear regression analysis to investigate the relationship between polygenic risk scores for SCZ and the thickness of RNFL, GCIPL, and INL. Because the UK Biobank data is prone to potential outliers and heteroscedasticity, we used robust regression, which employs M-estimation for robust linear modeling. M-estimators are a broad class of estimators that generalize maximum likelihood estimators, which are sensitive to outliers and violations of normality assumptions. The Huber weight function was applied to downweight the influence of outliers and heavy-tailed distributions on the estimation of model parameters[40]. The analyses were performed with the *rlm* function of the *MASS* package[41] in R, with the three retinal layer thickness measurements (averaged across both eyes) as the dependent variables and SCZ-PRS as the independent variable, while adjusting for age, age squared, genetic sex, body mass index, smoking status, hypertension, optical coherence tomography image quality, genotype array, and the first 10 genetic ancestry principal components.

### eFigures

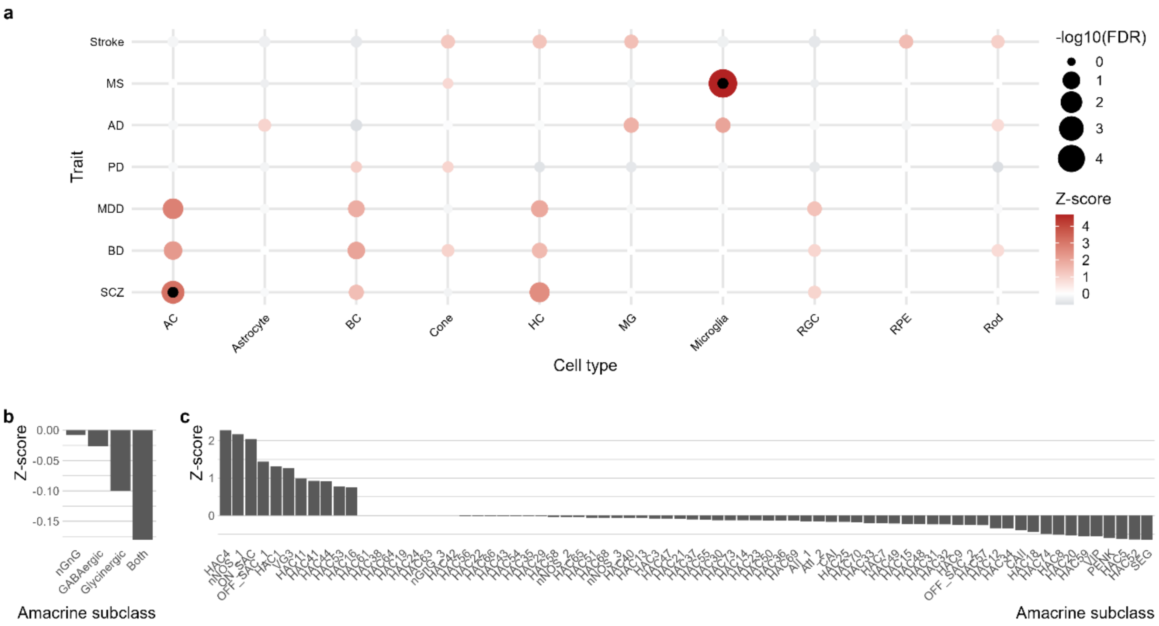


**eFigure 1: Association between genetic risk for schizophrenia and amacrine cells can be replicated in an independent human snRNA-seq data set and is not specific to a particular amacrine cell subtype.**

**a,** Dotplot illustrating the associations between retinal cell types and GWAS traits in the integrated Human Retinal Cell Atlas snRNA-seq data set, which comprises 1,775,529 nuclei[13]. Inner black dots indicate significant associations between amacrine cells and schizophrenia and between microglia and multiple sclerosis. The size of the outer dots represents the false discovery rate-adjusted MAGMA *P* value. The color scale illustrates the MAGMA Z-score. Significance was defined at a Bonferroni threshold of 0.5/7, corresponding to the seven traits under investigation. .

**b,** Bar plot illustrating enrichment of schizophrenia risk for major amacrine cell subtypes as shown by an additional MAGMA analysis restricted to the subset of 387,069 amacrine cells from the Human Retinal Cell Atlas[13]**.** No significant enrichment was found for GABAergic, glycinergic, both GABAergic and glycinergic, or non-GABAergic non-glycinergic subpopulations.

**c,** Barplot illustrating the enrichment between schizophrenia risk and 73 amacrine cell subtypes from the Human Retinal Cell Atlas snRNA-seq data set[13]. Analysis was restricted to the amacrine cell subset. None of the effects was statistically significant after FDR adjustment.

*Abbreviations:* AC, amacrine cell; AD, Alzheimer disease; BC, bipolar cell; BD, bipolar disorder; FDR, false discovery rate-adjusted *P* value; HC, horizontal cell; MDD, major depressive disorder; MG, Müller glia; MS; multiple sclerosis; nGnG, non-GABAergic non-glycinergic; PD, Parkinson disease; RGC, retinal ganglion cell; RPE, retinal pigment epithelium; SCZ, schizophrenia.

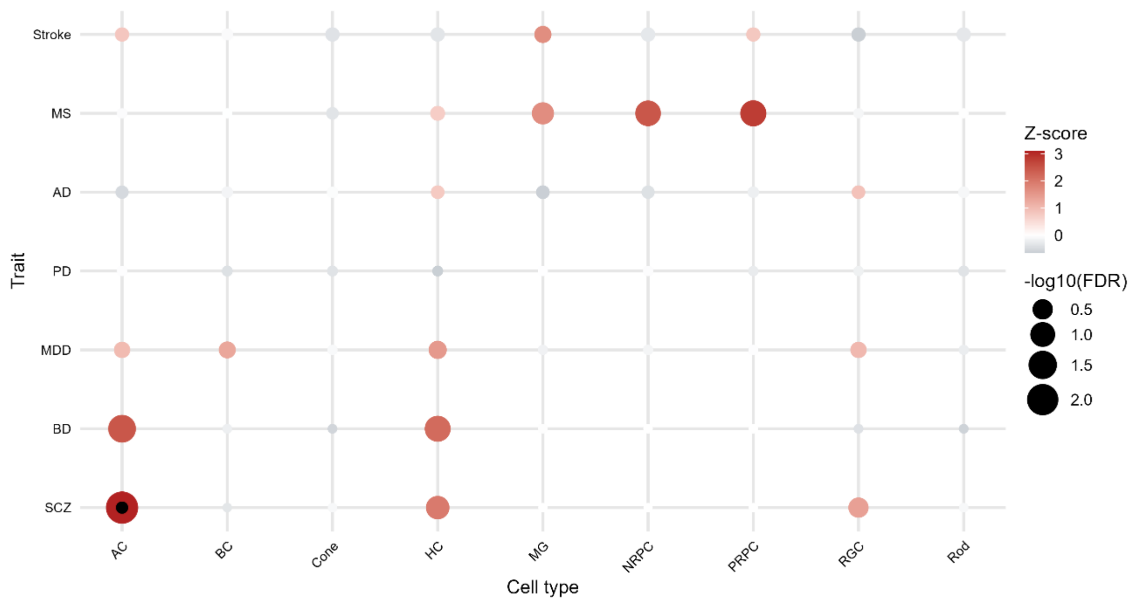


**eFigure 2: Schizophrenia polygenic risk converges on amacrine cells in the developing human retina.**

Dotplot illustrating the associations between cell types of fetal human retinas (9 to 24 weeks post fertilization)[14] and genome-wide association study summary statistics of major neurological and neuropsychiatric disorders. Inner black dots indicate statistical significance. The size of the outer dots represents the FDR-adjusted MAGMA *P* value. The color scale illustrates the MAGMA Z-score. *P* values were adjusted for the FDR within each trait, and the significance level was set to 0.05/7 after Bonferroni correction for the seven traits.

*Abbreviations:* AC, amacrine cell; AD, Alzheimer disease; BC, bipolar cell; BD, bipolar disorder; FDR, false discovery rat- adjusted *P* value; HC, horizontal cell; MDD, major depressive disorder; MG, Müller glia; MS; multiple sclerosis; NRPC, neurogenic retinal progenitor cell; PD, Parkinson disease; PRPC, primary retinal progenitor cell; RGC, retinal ganglion cell; RPE, retinal pigment epithelium; SCZ, schizophrenia.

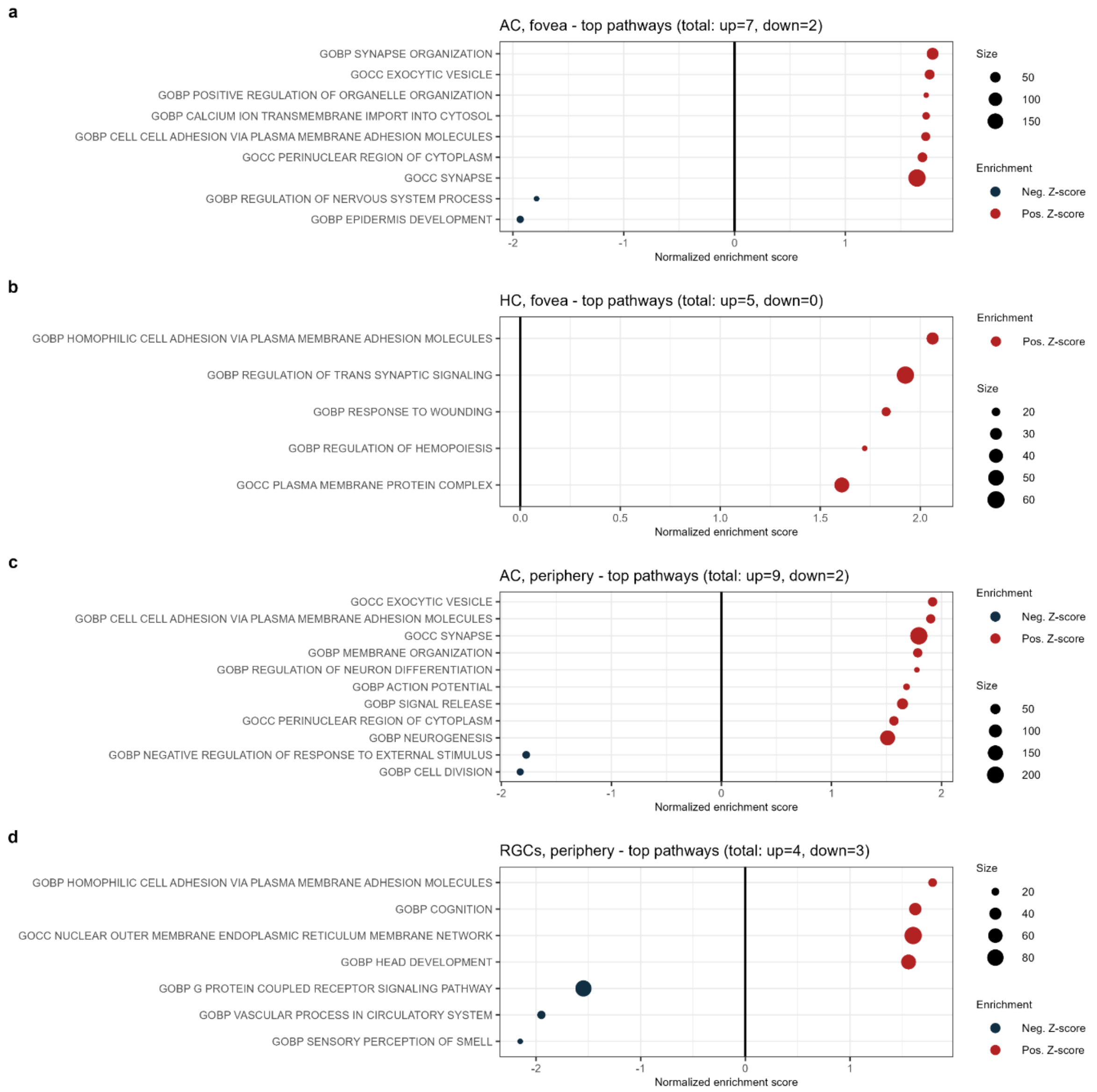


**eFigure 3: Gene set enrichment analysis points to altered synaptic and neuronal function in the retina in schizophrenia.**

Dotplots illustrate significant enrichment of Gene Ontology (GO) terms in those schizophrenia-associated genes that are specifically expressed in **a,** foveal amacrine cells, **b,** foveal horizontal cells, **c,** amacrine cells of the peripheral retina, and **d,** peripheral retinal ganglion cells, as calculated with a preranked gene set enrichment analysis. For the gene set enrichment analysis, the top 10% of genes with the highest expression specificity for each cell type in the human single-cell RNA sequencing data set of Cowan et al.[12] were included and genes were ranked by their MAGMA gene-level Z-scores for schizophrenia. Dot sizes represent the numbers of genes in the gene set overlapping with the top decile of cell type-specific genes for each cell type.

*Abbreviations:* AC, amacrine cell; GOBP, Gene Ontology biological process; GOCC, Gene Ontology cellular component; HC, horizontal cell; RGC, retinal ganglion cell.

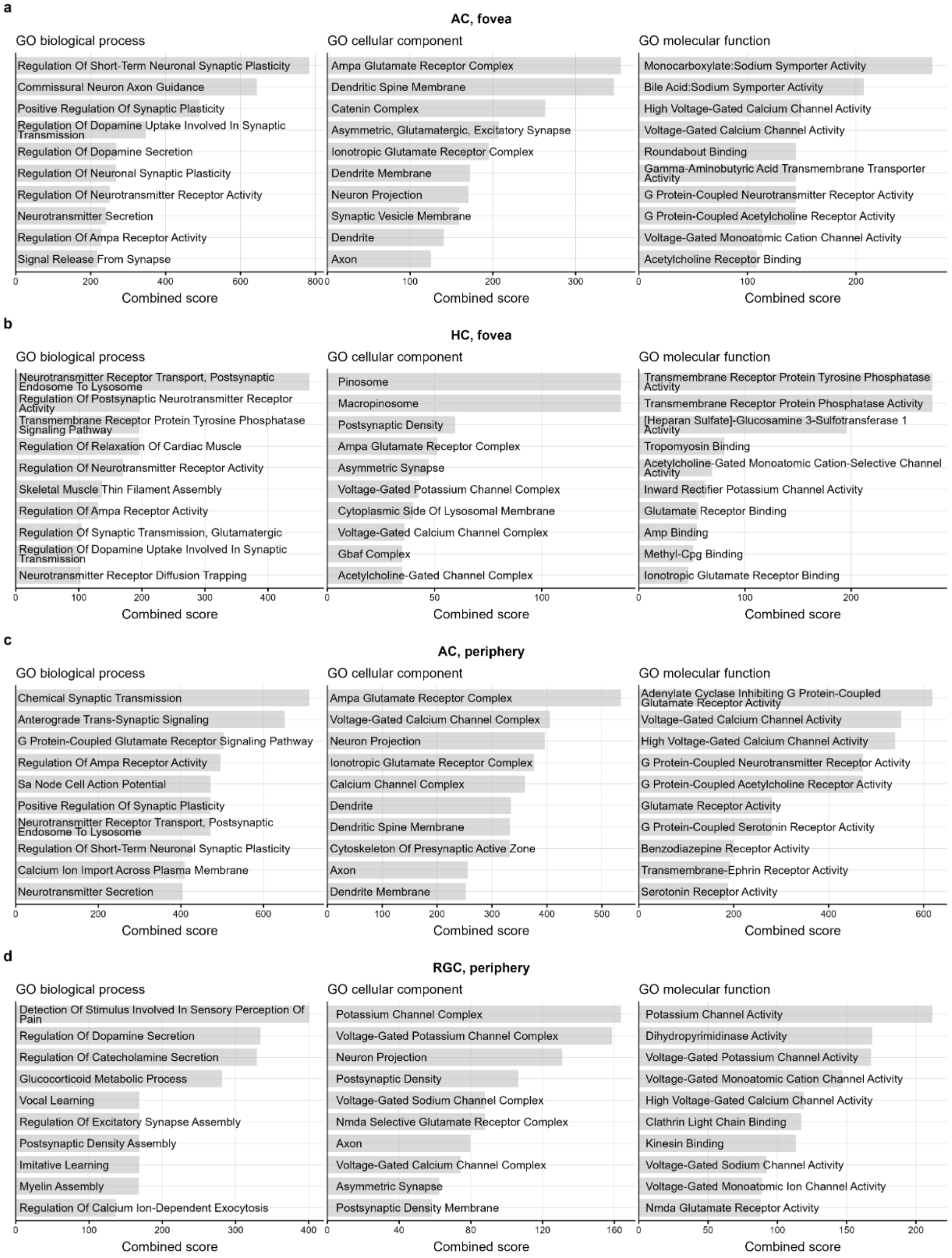


**eFigure 4: Enrichr cell type-specific Gene Ontology enrichment analysis for schizophrenia**

**a,** Bar plots illustrating the top 10 gene sets from Gene Ontology (GO) biological process (left), GO cellular component (middle), and GO molecular function (right) that were overrepresented among the 365 genes that had a MAGMA gene-level P value for schizophrenia (SCZ) less than 0.05 and were in the highest specificity decile for foveal amacrine cells. GO terms are ranked by the Enrichr combined score.

**b,** Bar plots illustrating the top 10 gene sets from GO biological process (left), GO cellular component (middle), and GO molecular function (right) that were overrepresented among the 330 genes that had a MAGMA gene-level *P* value for SCZ less than 0.05 and were in the highest specificity decile for foveal horizontal cells. GO terms are ranked by the Enrichr combined score.

**c,** Bar plots illustrating the top 10 gene sets from GO biological process (left), GO cellular component (middle), and GO molecular function (right) that were overrepresented among the 376 genes that had a MAGMA gene-level *P* value for SCZ less than 0.05 and were in the highest specificity decile for peripheral amacrine cells. GO terms are ranked by the Enrichr combined score.

**d,** Bar plots illustrating the top 10 gene sets from GO biological process (left), GO cellular component (middle), and GO molecular function (right) that were overrepresented among the 424 genes that had a MAGMA gene-level *P* value for SCZ less than 0.05 and were in the highest specificity decile for peripheral retinal ganglion cells. GO terms are ranked by the Enrichr combined score.

*Abbreviations:* AC, amacrine cell; HC, horizontal cell; GO, Gene Ontology; RGC, retinal ganglion cell.

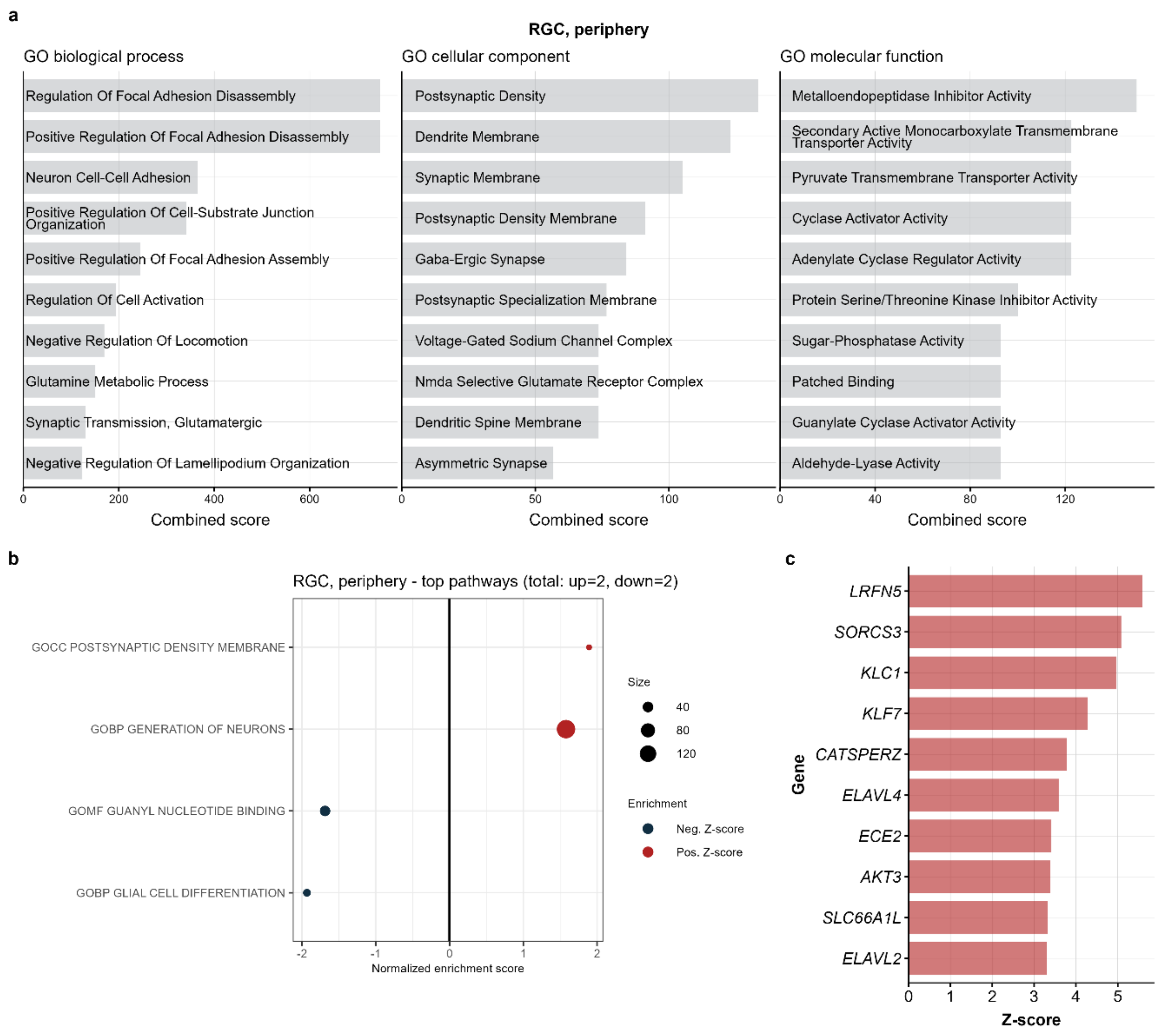


**eFigure 5: Gene Ontology enrichment analysis for major depressive disorder in peripheral retinal ganglion cells.**

**a,** Bar plots illustrating the 10 most enriched gene sets from Gene Ontology (GO) biological process (left), GO cellular component (middle), and GO molecular function (right) among the 138 genes that had a MAGMA gene-level *P* value less than 0.05 for major depressive disorder (MDD) and were within the highest decile of cell type specificity for human peripheral retinal ganglion cells, based on Enrichr. GO terms are ranked by the Enrichr combined score.

**b,** Preranked gene set enrichment analysis of GO terms in MDD-associated genes that are specifically expressed in peripheral retinal ganglion cells, based on MAGMA Z-scores for MDD. Dot sizes represent the numbers of genes in the gene set that overlapped with the top decile of cell type-specific genes for each cell type.

**c,** Bar plot indicating the top 10 MDD-associated genes with specific expression in peripheral retinal ganglion cells, ranked by their MAGMA gene-level Z-scores for MDD.

*Abbreviations:* GO, Gene Ontology; RGC, retinal ganglion cell.

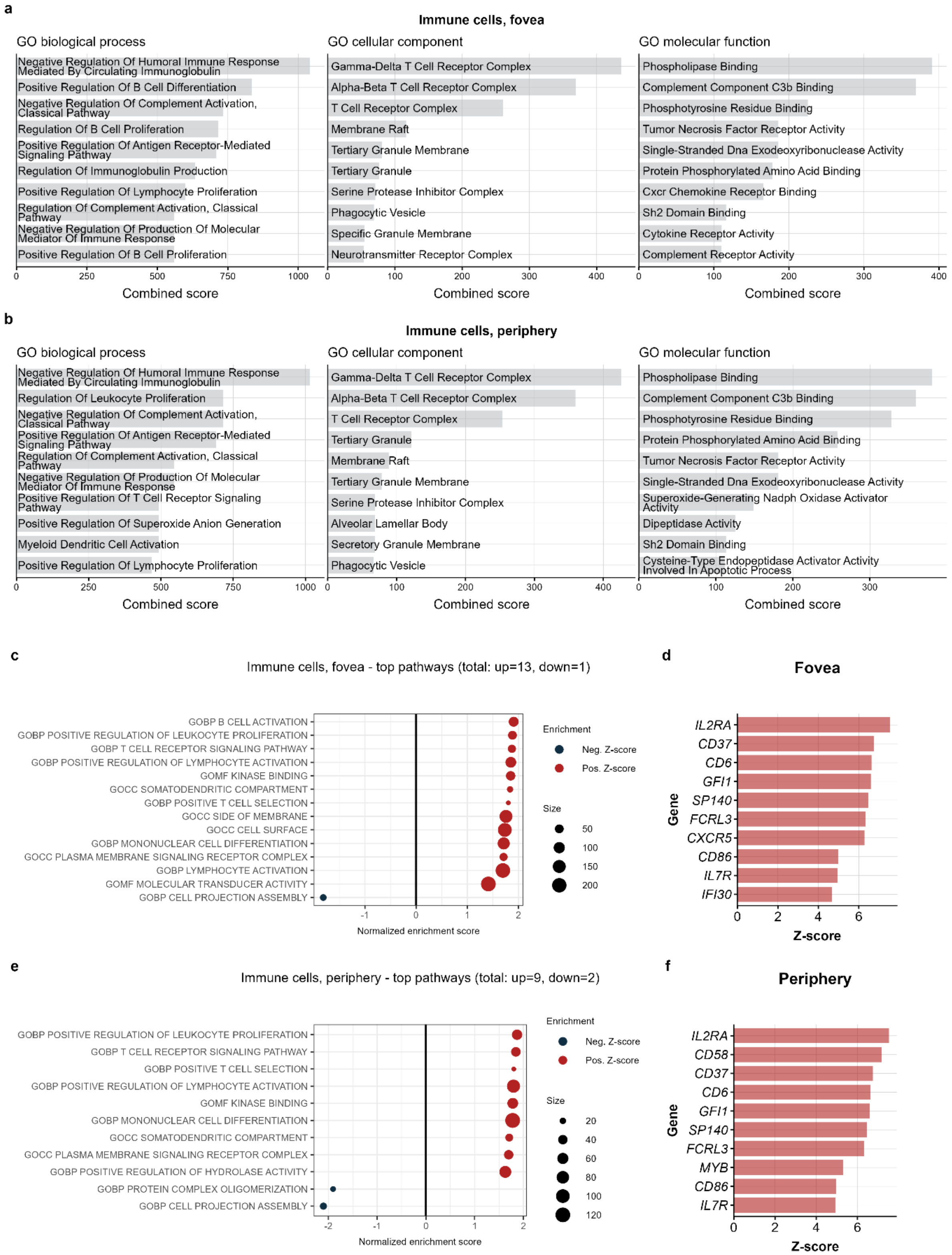
a
Immune cells, fovea
GO biological process
Negative Regulation Of Humoral Immune Response Mediated By Circulating Immunoglobulin
Positive Regulation Of B Cell Differentiation
Negative Regulation Of Complement Activation, Classical Pathway
Regulation Of B Cell Proliferation
Positive Regulation Of Antigen Receptor-Mediated Signaling Pathway
Regulation Of Immunoglobulin Production
Positive Regulation Of Lymphocyte Proliferation
Regulation Of Complement Activation, Classical Pathway
Negative Regulation Of Production Of Molecular Mediator Of Immune Response
Positive Regulation Of B Cell Proliferation
Combined score
GO cellular component
Gamma-Delta T Cell Receptor Complex
Alpha-Beta T Cell Receptor Complex
T Cell Receptor Complex
Membrane Raft
Tertiary Granule Membrane
Tertiary Granule
Serine Protease Inhibitor Complex
Phagocytic Vesicle
Specific Granule Membrane
Neurotransmitter Receptor Complex
Combined score
GO molecular function
Phospholipase Binding
Complement Component C3b Binding
Phosphotyrosine Residue Binding
Tumor Necrosis Factor Receptor Activity
Single-Stranded Dna Exodeoxyribonuclease Activity
Protein Phosphorylated Amino Acid Binding
Cxcr Chemokine Receptor Binding
Sh2 Domain Binding
Cytokine Receptor Activity
Complement Receptor Activity
Combined score
b
Immune cells, periphery
GO biological process
Negative Regulation Of Humoral Immune Response Mediated By Circulating Immunoglobulin
Regulation Of Leukocyte Proliferation
Negative Regulation Of Complement Activation, Classical Pathway
Positive Regulation Of Antigen Receptor-Mediated Signaling Pathway
Regulation Of Complement Activation, Classical Pathway
Negative Regulation Of Production Of Molecular Mediator Of Immune Response
Positive Regulation Of T Cell Receptor Signaling Pathway
Positive Regulation Of Superoxide Anion Generation
Myeloid Dendritic Cell Activation
Positive Regulation Of Lymphocyte Proliferation
Combined score
GO cellular component
Gamma-Delta T Cell Receptor Complex
Alpha-Beta T Cell Receptor Complex
T Cell Receptor Complex
Tertiary Granule
Membrane Raft
Tertiary Granule Membrane
Serine Protease Inhibitor Complex
Alveolar Lamellar Body
Secretory Granule Membrane
Phagocytic Vesicle
Combined score
GO molecular function
Phospholipase Binding
Complement Component C3b Binding
Phosphotyrosine Residue Binding
Protein Phosphorylated Amino Acid Binding
Tumor Necrosis Factor Receptor Activity
Single-Stranded Dna Exodeoxyribonuclease Activity
Superoxide-Generating Nadph Oxidase Activator Activity
Dipeptidase Activity
Sh2 Domain Binding
Cysteine-Type Endopeptidase Activator Activity Involved In Apoptotic Process
Combined score
c
Immune cells, fovea - top pathways (total: up=13, down=1)
GOBP B CELL ACTIVATION
GOBP POSITIVE REGULATION OF LEUKOCYTE PROLIFERATION
GOBP T CELL RECEPTOR SIGNALING PATHWAY
GOBP POSITIVE REGULATION OF LYMPHOCYTE ACTIVATION
GOMF KINASE BINDING
GOCC SOMATODENDRITIC COMPARTMENT
GOBP POSITIVE T CELL SELECTION
GOCC SIDE OF MEMBRANE
GOCC CELL SURFACE
GOBP MONONUCLEAR CELL DIFFERENTIATION
GOCC PLASMA MEMBRANE SIGNALING RECEPTOR COMPLEX
GOBP LYMPHOCYTE ACTIVATION
GOMF MOLECULAR TRANSDUCER ACTIVITY
GOBP CELL PROJECTION ASSEMBLY
Normalized enrichment score
Enrichment
Neg. Z-score
Pos. Z-score
Size
50
100
150
200
d
Fovea
IL2RA
CD37
CD6
GFI1
SP140
FCRL3
CXCR5
CD86
IL7R
IFI30
Gene
Z-score
e
Immune cells, periphery - top pathways (total: up=9, down=2)
GOBP POSITIVE REGULATION OF LEUKOCYTE PROLIFERATION
GOBP T CELL RECEPTOR SIGNALING PATHWAY
GOBP POSITIVE T CELL SELECTION
GOBP POSITIVE REGULATION OF LYMPHOCYTE ACTIVATION
GOMF KINASE BINDING
GOBP MONONUCLEAR CELL DIFFERENTIATION
GOCC SOMATODENDRITIC COMPARTMENT
GOCC PLASMA MEMBRANE SIGNALING RECEPTOR COMPLEX
GOBP POSITIVE REGULATION OF HYDROLASE ACTIVITY
GOBP PROTEIN COMPLEX OLIGOMERIZATION
GOBP CELL PROJECTION ASSEMBLY
Normalized enrichment score
Enrichment
Neg. Z-score
Pos. Z-score
Size
20
40
60
80
100
120
f
Periphery
IL2RA
CD58
CD37
CD6
GFI1
SP140
FCRL3
MYB
CD86
IL7R
Gene
Z-score

**eFigure 6: Enrichment analysis highlights immunological Gene Ontologies in multiple sclerosis.**

**a**–b, Bar plots illustrating the top 10 most enriched gene sets from Gene Ontology (GO) biological process (left), GO cellular component (middle), and GO molecular function (right) in the lists of genes that were significantly associated with multiple sclerosis (MS) and specifically expressed in retinal immune cells of the fovea (**a**) or the peripheral retina (**b**). Analysis was performed with Enrichr and based on the 209 (fovea) and 213 (peripheral retina) genes with specific expression in the immunce cell population and a MAGMA gene-level *P* value for MS less than 0.05. Gene sets are ranked by the Enrichr combined score.

**c,** Dotplot illustrating the enrichment between GO terms and MS within those genes that were specifically expressed in foveal immune cells, as obtained by a preranked gene set enrichment analysis (GSEA). For the GSEA, genes were ranked by their MAGMA Z-score for MS.

**d,** Bar plot indicating the top 10 MS-associated genes with specific expression in foveal immune cells, ranked by their MAGMA gene-level Z-scores for MS.

**e,** Dotplot illustrating the enrichment between GO terms and MS within those genes that were specifically expressed in peripheral retinal immune cells, as obtained by a preranked GSEA. For the GSEA, genes were ranked by their MAGMA Z-score for MS.

**f,** Bar plot indicating the top 10 MS-associated genes with specific expression in peripheral retinal immune cells, ranked by their MAGMA gene-level Z-scores for MS.

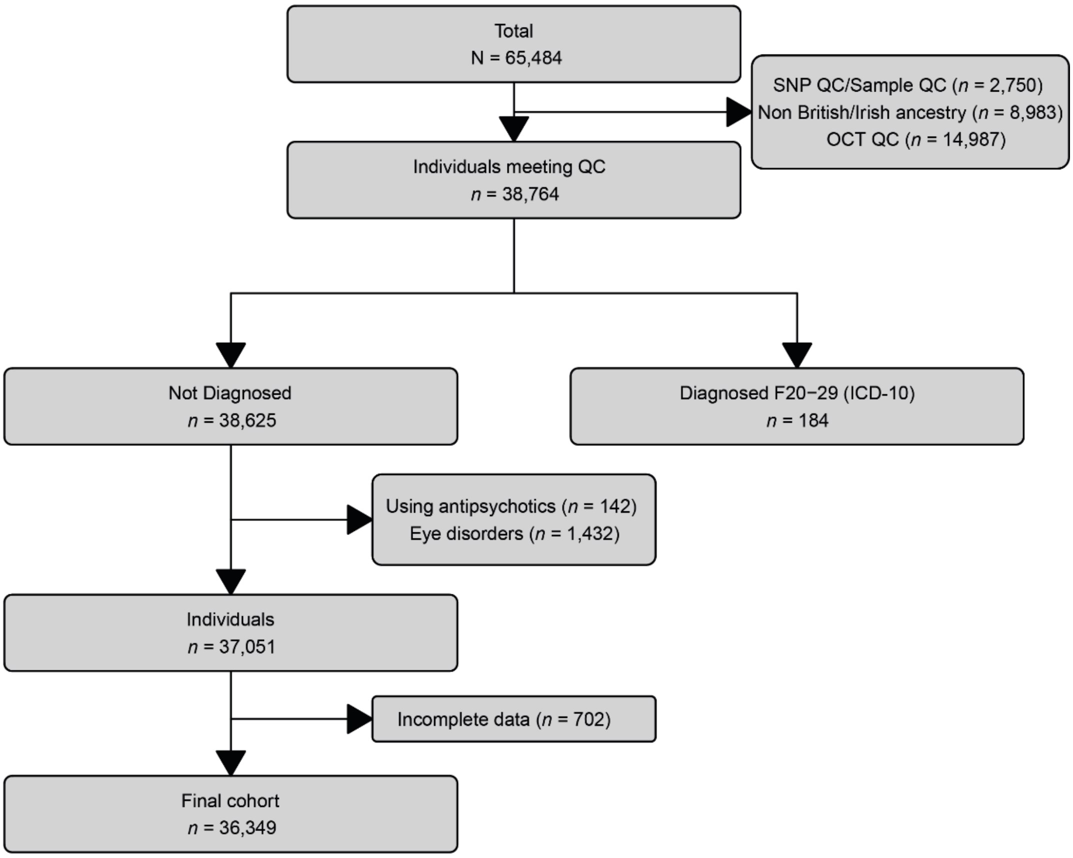


**eFigure 7: Flowchart highlighting the inclusion and exclusion criteria and showing how the final UK Biobank cohort was identified.**

*Abbreviations:* ICD-10, 10th revision of the International Statistical Classification of Diseases and Related Health Problems; OCT, optical coherence tomography; QC, quality control; SNP, single nucleotide polymorphism.
